# Natural Zeitgebers cannot compensate for the loss of a functional circadian clock in timing of a vital behaviour in *Drosophila*

**DOI:** 10.1101/2019.12.22.886309

**Authors:** Franziska Ruf, Oliver Mitesser, Simon Tii Mungwa, Melanie Horn, Dirk Rieger, Thomas Hovestadt, Christian Wegener

## Abstract

The adaptive significance of adjusting behavioural activities to the right time of the day is intuitive. Laboratory studies have implicated an important role of circadian clocks in behavioural timing and rhythmicity. Yet, recent studies on clock-mutant animals questioned this importance under more naturalistic settings, as various clock mutants showed nearly normal diel activity rhythms under semi-natural Zeitgeber conditions.

We here report evidence that proper timing of eclosion, a vital behaviour of the fruit fly *Drosophila melanogaster*, requires a functional molecular clock even under quasi-natural conditions. In contrast to wildtype flies, *period^01^* mutants with a defective molecular clock eclose mostly arrhythmically in a temperate environment even in the presence of a full complement of abiotic Zeitgebers. Moreover, *period^01^* mutants eclose during a much larger portion of the day, and peak eclosion time becomes more susceptible to variable day-to-day changes of light and temperature. Under the same conditions, flies with impaired peptidergic inter-clock signalling (*pdf^01^* and *han^5304^* mutants) stayed largely rhythmic with normal gate sizes. Our results suggest that the presence of natural Zeitgebers can mitigate a loss of peptide-mediated phasing between central clock neuron groups, but cannot substitute for the lack of a functional molecular clock under natural temperate conditions.

## Background

Endogenous timing via circadian clocks confers adaptive advantages as it allows organisms to anticipate daily changes in the environment (see [1–3]). In terms of behaviour, the fitness relevance of being able to schedule locomotor activity, feeding, mating or other actions at the right time of the day is intuitive as it may help maximize success and reduce risks. Many studies under constant laboratory conditions have revealed a key role of the central and peripheral clocks in timing of behaviours across taxa. However, the importance of circadian clocks in daily timing of behaviours under natural conditions or in ecological context has come under debate, as studies in the last decade have assessed the functional importance of endogenous clocks under (semi-) natural conditions in a variety of mostly vertebrate species (see [1,2,4]). One important conclusion derived from these studies is that diel activity rhythms can remarkably differ between seminatural and laboratory conditions, since the phase relationship between behavioural activity and a given Zeitgeber such as light is modulated by other abiotic Zeitgebers, particularly by temperature [5]. Furthermore, intra- and interspecific interactions such as predation [6–8] or competition for food [9] determine (“mask”) activity patterns in the wild. Most strikingly, under semi-natural conditions in an outdoor enclosure, *Per2^BRDM1^*mice carrying a mutation in a core clock gene showed the same activity pattern as controls, and both showed mostly diurnal feeding, although they are strongly nocturnal in laboratory conditions [10]. In the wild, chipmunks with a lesion in the suprachiasmatic nucleus (SCN, the “master clock” in mammals) showed above-ground activity pattern similar to controls, even though SCN lesioned animals exhibit more night-time activity inside the den [11,12]. In case of voles and larger mammals, feeding and energy metabolism appear to be key drivers of diel activity patterns [9,13] that mask circadian control. Together, these findings appear to challenge “circadian dominance” [13], and instead attribute more importance to external and internal Zeitgebers in daily timing of behaviours in nature.

The fruit fly (*Drosophila melanogaster*) is the best studied invertebrate model in circadian research. *Drosophila* shows robust behavioural rhythms in the laboratory, including locomotor activity (see [14]). Under normal light:dark (LD) conditions, *Drosophila* is characterised by a biphasic locomotor activity pattern with a morning (M) and evening (E) activity peak around lights-on and lights-off [15–17]. This pattern is maintained in constant darkness (DD) and has been highly reproducible. It came thus as a surprise when the first study using activity monitors placed outdoors reported that fruit flies behave quite differently under quasi-natural conditions, becoming more diurnal than crepuscular [18]. At higher temperatures, the flies exhibited increased midday locomotor activity (A peak) instead of the siesta phase seen in the laboratory even under similarly high temperatures. While demonstrating strongly impaired rhythmicity under light:dark conditions and arrhythmicity under constant conditions in the laboratory (e.g. [15,19], clock mutants and flies with genetically ablated circadian pacemaker cells showed a high degree of locomotor rhythmicity with little difference in activity patterns compared to wildtype controls [18], reminiscent of the results for *Per2^BRDM1^* mice [10]. The onset of morning activity in wildtype and clock-impaired flies was inversely related with night temperature and hence seems to represent a temperature response rather than a clock-controlled activity. The onset of the evening activity peak was in contrast clock-dependent at lower temperatures, with a strong temperature modulation at higher temperatures [18]. In the laboratory, nature-like simulated twilight regime is sufficient to induce wildtype-like locomotor rhythmicity with M and E peak activity in *per^01^* and *tim^01^* clock mutants even at constant temperature [20], providing further evidence for a subordinate role of the circadian clock in controlling the daily locomotor activity pattern. While these studies question the “circadian dominance” of daily activity patterns, they all are restricted to locomotor activity which is a component common to a multitude of ecologically important behaviours in nature, from foraging and social interactions to escape from predators or adverse abiotic conditions. Locomotor activity is thus a behavior sensitive to a variety of environmental stimuli, and the endogenous locomotor activity rhythms are prone to masking.

We therefore hypothesised that a “circadian dominance” with all its intuitive and likely advantages [2–4] may become more evident in other behaviours, especially those that serve only one particular function. Eclosion, the emergence of the adult holometabolic insect from the pupa, is arguably one of the most specific behaviours in insects, with only one dedicated function (propelling the pharate adult out of the pupal case). Though it occurs only once in a lifetime, in *Drosophila* it is a rhythmical event on the population level gated to dawn by the interaction of the central clock in the brain and a peripheral clock in the steroid-producing prothoracic gland [21–23]. Moreover, once initiated, eclosion behaviour cannot be interrupted and stereotypically follows a fixed action pattern [24–27], a condition under which internal timing might be especially important. Unlike locomotor and most other behaviours, eclosion is basically free of motivational states and inter-individual interactions; flies start to feed, mate and interact only several hours after eclosion. Eclosion rhythmicity of *Drosophila* can be entrained by light and temperature changes [28], and once entrained rhythmicity is stable under constant conditions even if the fly population was synchronised by only a brief light pulse or temperature step during larval development [28–31]. Flies with a mutation in the core clock genes *period* (*per*) or *timeless* (*tim*) eclose arrhythmically not only under constant conditions but also under laboratory LD cycles [28,32–34], emphasizing the requirement of a functioning molecular clockwork for eclosion rhythmicity.

We here report our results on the eclosion rhythmicity of wildtype and clock mutant flies under quasi-natural temperate conditions using a newly developed open eclosion monitor [35]. Our results show that a functional molecular clock is required for behavioural rhythmicity under these conditions. While natural Zeitgebers can compensate a loss of peptide-mediated phasing between central clock neuron groups, they are unable to substitute a functional molecular clock under temperate conditions.

## Methods

### Flies

For the eclosion experiments, the following fly strains were used: wildtype Canton S (WT_CS_), *w^+^ per^01^*[36], *w^+^ per^01^;tim^01^*, *w^+^ han^5304^*[37] and *w^+^;;pdf^01^* [38] (kind gifts of Charlotte Förster (Würzburg, Germany)). The *pdf^01^* line had been cantonised by Taishi Yoshii a few years ago; the genetic backgrounds of the other mutants are not equivalent. Flies were raised on standard Drosophila food medium (0.8% agar, 2.2% sugarbeet syrup, 8.0% malt extract, 1.8% yeast, 1.0% soy flour, 8.0% corn flour and 0.3% hydroxybenzoic acid).

### Eclosion under laboratory conditions

In the laboratory, flies were entrained either under 12h light, 12h dark (LD 12:12) regime at 20°C and 65% humidity (light entrainment), or at constant red light (λ =635 nm) and 65% humidity with 12hours at 25°C, 12h at 16°C (WC 25:16, temperature entrainment), or at constant red light (λ =635 nm) at 20°C and 12h at 70% humidity, 12h at 30% relative humidity (HD70:30; humidity entrainment). For WC entrainment, temperature was ramped by 0.1°C/min between conditions. Eclosion was monitored by the open WEclMon system [35]. Puparia were individually taken out of the culture vials and glued onto a platform on eclosion plates with a cellulose-based glue (Tapetenkleister No. 389; 1:30, Auro, Germany). At the end of the light phase of day 0, eclosion plates were mounted in the WEclMons and eclosion was monitored for one week at 20°C, either under LD 12:12 or WC25:16 at 65% humidity. Infrared light (λ=850 nm) was given throughout the experiment.

To assess the effect of relative humidity on eclosion success, flies were allowed to lay eggs over night on standard food petri dishes in big egg-laying vials with mesh on top. Afterwards, adult flies were removed and the eggs/larvae were kept in a climate chamber (25 °C ± 0.2 °C; 60 % ± 2 % rH) under LD 12:12. After pupariation, the food source was removed and the meshvials were transferred to an incubator set at different relative humidity values (2 %, 60 % or 80 %) but otherwise similar conditions (LD 12:12, 25 °C ± 0.2 °C) one or two days prior to eclosion. The light regime and temperature inside the incubator was kept the same as during development. After a few days, successfully eclosed flies as well as unopened puparia were counted, and eclosed flies were checked for successful wing expansion.

### Eclosion under natural conditions

Experiments under natural conditions were conducted from July to October 2014 and July to September 2016 in a shelter shaded by bushes at the bee station/Hubland campus of the University of Würzburg (49° 47’ N, 9° 56’ E) [35]. The shelter was roofed and open at three sides. To keep predatory insects out and to prevent flies from escaping to the environment, the open sides were stretched with air-permeable black gauze. Double-sided sticky tape was glued around each monitor to trap predators and freshly eclosed flies. Flies were continuously bred inside the shelter in large 165 ml plastic culture vials (K-TK; Retzstadt, Germany), and were allowed to lay eggs for 3 to 4 days per vial to provide puparia in different developmental stages. At least six vials were kept in parallel. Once most of the larvae had pupariated, vials were transferred to the laboratory and puparia were collected and fixed on eclosion plates as described above. The loaded eclosion plates were then directly placed back into the shelter and eclosion was monitored for one week under constant red (λmax=635 nm) or infrared (λmax=850nm) light using the open Würzburg Eclosion Monitors (WEclMon [35]). Eclosion rhythmicity was found to be similar under red or infrared illumination [35]. Light intensity, temperature and relative humidity were registered with a MSR 145S datalogger (MSR Electronics GmbH, Seuzach, Switzerland), placed directly at the side of the monitors.

### Data analysis

Rhythmicity and period length of the eclosion profiles were analyzed by a toolbox developed by Joel Levine [39] in MATLAB (MathWorks, Inc., Natick, USA). A rhythmicity index (RI) was calculated by autocorrelation analysis. An RI > 0.3 indicates strongly rhythmic behaviour, 0.1<RI<0.3 indicates weakly rhythmic behaviour, while an RI<0.1 indicates arrhythmicity.

Statistical analysis with one-way ANOVA followed by Tukey post-hoc test and independent-samples t-test were performed with R (version 3.2.0; https://www.r-project.org/). Circular-linear correlation and circular statistics (mean, vector length, standard deviation) were analysed and plotted using Oriana 4.02 (Kovach Computing Services, Pentraeth, Isle of Anglesey, UK). Graphs of outdoor eclosion profiles were compiled in R (version 3.2.0).

### Logistic regression model

We analyzed the data for the day-time of eclosion using a logistic regression approach and thestatistics software R (https://www.r-project.org/). The focus of our analysis was the timing of eclosion within a day whereas the day (date) of emergence is treated as a random factor. Hour of day was included as a linear and a quadratic predictor. Relative humidity and temperature were strongly correlated (R=0.9, p<0.001), and for reasons below we decided to exclude humidity from the analysis. To avoid computational problems and allow for comparison of the impact of different factors we standardized variables hour of day (centered around 12:00), temperature, light intensity, and nautical dawn (data provided by www.timeanddate.de) by centering with respect to their root mean square. We further only included data for days 2-5 of each experiment as too few flies eclosed on the other days of the experiments. The R library “GLMMadaptive” version 0.6-5 [40] (Rizopoulos 2019, vers. 0.6-5) was used for model fitting with the fixed explanatory variables genotype (factorial), hour and hour^2^, temperature, light intensity and nautical dawn nested into the random factor “experimental group” and as dependent variable the proportion of flies emerging within a given daily hour out of those flies that had not yet emerged before on this day. We assumed that flies emerging on different days were emerging from eggs laid on different days and ignored interactions of higher than second order; the interaction term between hour and hour^2^ was not included in the model. Our main interest was in understanding the temporal emergence pattern over the course of day.

We selected the best statistical model by backward simplification as suggested by Crawley [41], starting with the most complex model and removing effects of weak significance until Anova model comparison yielded a significant difference indicating that further simplification would result in a substantially worsening of the model.

## Results and Discussion

### Eclosion rhythmicity under laboratory light and temperature entrainment in WT_CS_ and *per^01^* clock mutants

As eclosion rhythmicity can be entrained by light and temperature, we first characterised the temporal eclosion pattern of wildtype Canton S (WT_CS_) and *per^01^* clock mutant flies under 12 h light:12h dark (LD12:12) and 12h warm:cold (WC12:12) light or temperature entrainment. WT_CS_ control flies were rhythmic under LD12:12 (Fig. 1A) and maintained rhythmicity in DD after light entrainment (Fig. 1A”) while *per^01^* mutants eclosed arrhythmically under both conditions (Fig. 1B, B”). The WT_CS_ showed the typical daily eclosion profile with bouts of eclosion events during a gate in the hours around ZT0 (lights on), with strongly reduced or absent eclosion activity in the second half of the day (ZT12-23). Under temperature entrainment, WT_CS_ eclosed rhythmically (Fig. 1A’), but this rhythmicity dampened strongly in CC (Fig. 1A’”). The *per^01^* mutants showed weak diel rhythmicity under WC conditions with a strong ultradian component with a period length of around 12h (Fig. 1B’), yet turned arrhythmic in CC (Fig. 1B’”). These results are largely consistent with previous studies [28,33,42] and indicate that a functional clock is required to drive normal eclosion rhythmicity under LD and WC as well as constant conditions.

**Figure 1:**
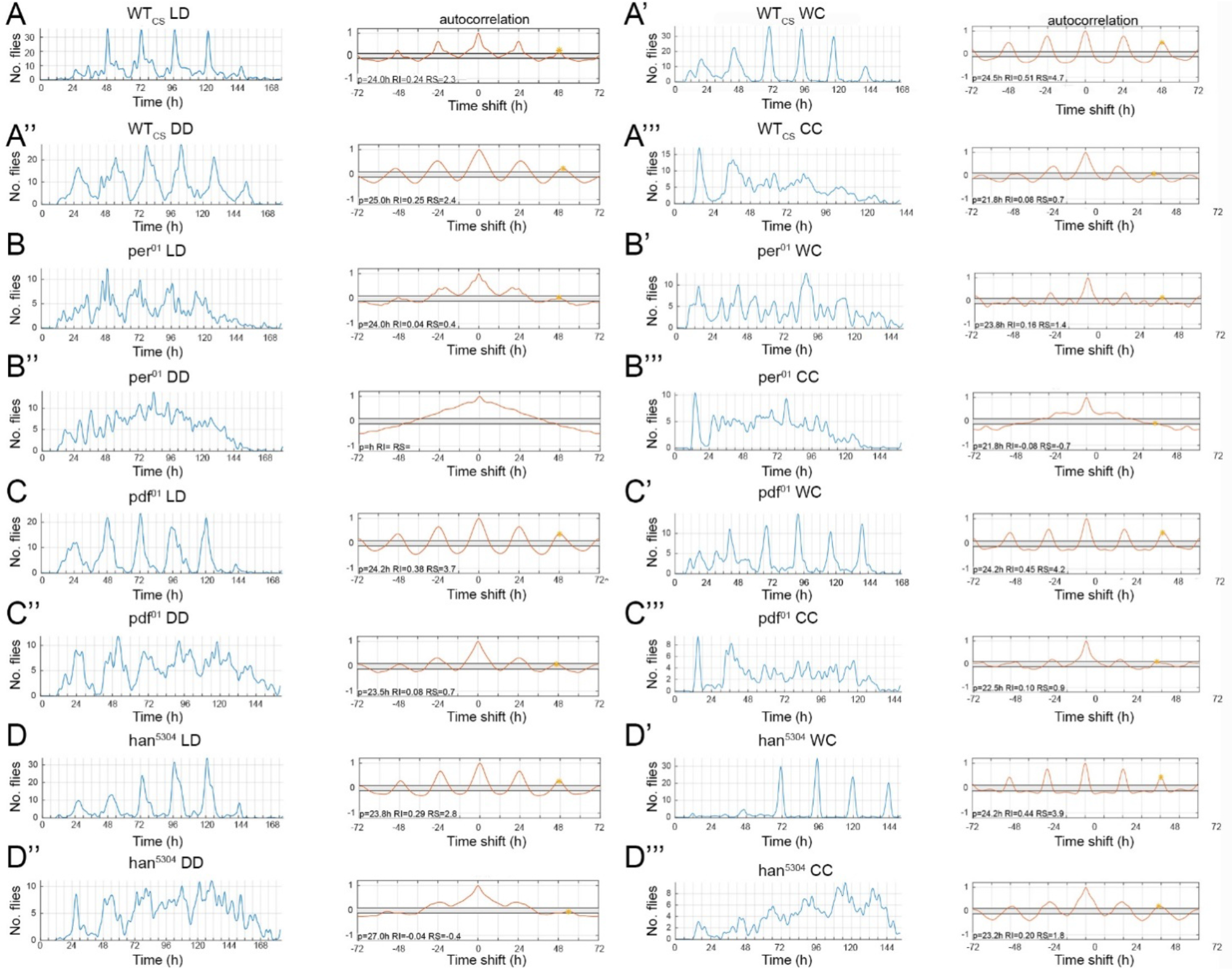
Eclosion profiles and autocorrelation analysis of WT_CS_, per^01^ and PDF signalling mutants under laboratory conditions. Left half: light entrainment (LD 12:12), right half: temperature entrainment (WC 25:16). **A-A’**) WT_CS_ flies eclose rhythmically under ZT conditions as well as under constant darkness (DD) yet quickly loose rhythmicity under constant temperature (**A’ lower panel**). In contrast, per^01^ clock mutants show impaired rhythmicity under all conditions (**row 3**: ZT conditions, **row 4**: constant conditions). Under WC entrainment, however, ultradian rhythmicity appeared with a period of 12h. Flies lacking either PDF (**row 5**) or the PDF receptor (han^5304^, **row 7**) eclose rhythmically during ZT conditions, but increasingly lose rhythmicity during constant conditions (**row 6 and 8**).

### Weaker eclosion rhythmicity under quasi-natural conditions in wildtype flies

To test for eclosion rhythmicity under natural conditions, we assayed eclosion rhythmicity outdoors, using the open WEclMon system. Pupariae were directly exposed to the ambient changes in temperature, humidity and indirect changes in light intensity without interfering plastic or glass interfaces as used by previous studies under semi-natural conditions [43,44]. Under these quasi-natural conditions, WT_CS_ flies eclosed rhythmically in the majority of experiments (78%, n=9, Fig. 2A’), as was expected from flies with intact circadian clock receiving continuous Zeitgeber information. The proportion of rhythmic experiments was similar compared to flies raised and monitored under light-dark (LD12:12) conditions in the laboratory (75%of rhythmic eclosion (n=4, Fig. 2A’)). Yet, robustness of eclosion rhythms of WT_CS_ flies under quasi-natural conditions was always weak (rhythmicity index 0.1<RI<0.3), while at least one out of four experiments under LD12:12 showed very robust eclosion rhythm (RI > 0.3, Fig. 2A’). The mean daily hour for eclosion over all WT_CS_ experiments was 09:06 h ± 04:23 s.d. (n =5730), with a mean vector length (r) of 0.517 (Fig. 2A’”) as obtained by circular statistics. Under laboratory warm-cold conditions (WC12:12) alternating between 25°C and 16°C, proportion of rhythmic experiments was increased to 92%, with five out of 12 experiments showing strong rhythmicity (Fig. 2A’). Compared to quasi-natural and LD conditions, the rhythmicity was slightly but not significantly increased under WC conditions (Fig. 2A”) suggesting that stable temperature cycles promote robust eclosion rhythms. In line, the strongest rhythm under quasi-natural conditions occurred when amplitude and shape of temperature oscillations remained stable over the course of experiment (6 days) and stayed within the range of approximately 15-30°C (Suppl. Fig. 1A).

**Figure 2:**
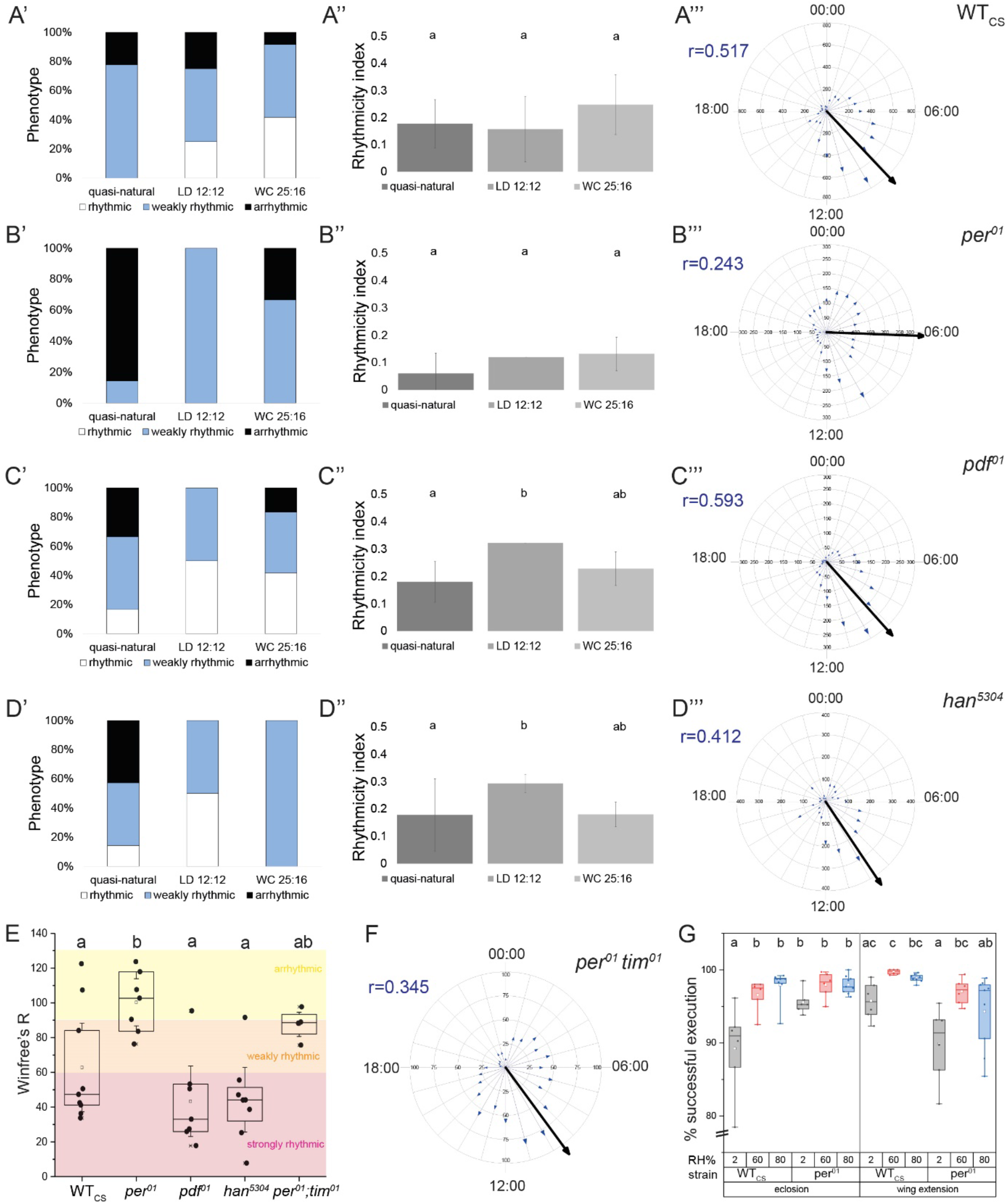
*Eclosion rhythmicity under quasi-natural conditions in **A)** WT_CS_, **B)** per^01^, **C)** pdf^01^ **D)** han^5304^ and **F)** per^01^, tim^01^ flies and **G)** successful eclosion and post-eclosion of WT_CS_ under varying relative humidity. The left column **(A′-D′)** summarises the rhythmicity of individual experiments under quasi-natural conditions, and in LD12:12 and WC25:12 entrainment in the laboratory. The middle column **(A″-D″)** shows the mean rhythmicity index (RI) ± s.d. for the different conditions*. The right column (***A′″-D′″*)** shows circular plots of the data with mean vector (black arrow) and sum of eclosed flies per hour over all experiments (blue arrows). *E) Winfree’s rhythmicity index for the different genotypes under quasi-natural conditions, **F)** Results of the circular analysis with the mean eclosion time (black arrow) of the length r as indicated. The blue arrows give the number of eclosions at a given time. **G)** Percentage of flies that either successfully eclosed (left) or extended wings among those that did eclose (right). Small letters indicate statistical significance (p<0.05). A’ has already been published in a different context [35]*.

High temperatures (max. 33°C) and mild nights above 20°C had little effect on peak eclosion time (ψ_PK_), while low night temperatures < 15°C seemed to shift eclosion later into the day (Suppl. Fig. 1B). This is reflected by a stronger correlation for minimum day temperature and ψ_PK_ (r=0.383, p<0.001), while the circular-linear correlation between maximum day temperature and peak eclosion time was only weak (ψ_PK_, r=0.212, p=0.056). In both laboratory [31,42] and semi-natural [44] experiments under tropical conditions, ψ_PK_ delayed with decreasing temperature, in line with our results. Yet, ψ_PK_ advanced with increasing temperature [31,42,44] which was not observed in our experiments under temperate conditions.

### *per^01^* flies loose eclosion rhythmicity under quasi-natural conditions

Under quasi-natural temperate conditions,*per^01^* flies eclosed arrhythmically (RI<0.1) in the majority of experiments (86%, n=7, Fig. 2B’), even though these flies received continuous Zeitgeber information. As a consequence, the mean RI under quasi-natural conditions was markedly though not significantly reduced compared to laboratory conditions (Fig. 2B”). This finding is in strong contrast to observations under tropical semi-natural conditions where eclosion rhythmicity of *per^01^* flies was higher compared to laboratory conditions [43]. Importantly, the time window in which most of the eclosion events occurred (the eclosion gate) was considerably wider when compared to WT_CS_ flies; eclosion events occurred more frequently especially during the second half of the night (Fig. 2B’”). Accordingly, the mean eclosion time was around 3h earlier than in WT_CS_ flies with considerably high variance (06:08 h ± 06:25 s.d. (n =2449), r=0.243 (Fig. 2B’”)) as expected from flies that do not have to wait for the circadian eclosion gate to open, but can directly eclose after maturation. Nonetheless, even in *per^01^* flies, eclosion appears to be weakly gated with hardly any eclosion in the afternoon similar to that in WT_CS_ where the “forbidden” phase is extended into the first half of the night (Fig. 2A’”).

To test for eclosion gate width, we calculated Winfree’s R [45] which is defined as the number of eclosions outside an 8-hour gate, divided by the number of eclosions within this gate, multiplied by 100. R < 60 represents strong eclosion rhythmicity, 60< R< 90 represents weak rhythmicity, and R >90 represents arrhythmic eclosion (see [46]). Based on Winfree’s R, 67% of the quasi-natural experiments for WT_CS_ flies were strongly rhythmic, 11% were weakly rhythmic, and 22% were arrhythmic (n=9). None of the quasi-natural experiments for *per^01^* mutant flies showed strong rhythmicity, 29% were weakly rhythmic and 71% were arrhythmic (n=7). The mean R of *per^01^* mutant flies was 100.3 ± 6.7 s.d., and significantly higher (p=0.04, Tukey Multiple comparisons) than in WT_CS_ (62.8 ± 11.1 s.d., Fig 2E).

Winfree’s R for our laboratory data gave weak eclosion rhythmicity under LD12:12 (n=2), and mostly weak rhythmicity (n=6) under WC25:16 conditions for *per^01^* flies, suggesting that stable Zeitgeber amplitudes can narrow the eclosion gate and drive eclosion rhythmicity in flies with impaired clock. This would explain the finding of rhythmic eclosion of *per^01^* flies under tropical conditions [43], and is also in line with the finding that our sole rhythmic quasi-natural experiment in *per^01^* mutants occurred under stable temperature amplitudes with temperate highs (Suppl. Fig. 1A’). When Zeitgeber changes were shallow and did not show significant daily changes, eclosion became arrhythmic in *per^01^* mutant flies (Suppl. Fig. 1B’), while WTCs controls maintained rhythmicity (Suppl. Fig. 1B). Compared to WT_CS_ flies, *per^01^* mutants showed a stronger circular-linear correlation between maximum day temperature and ψ_PK_ (r=0.335, p=0.003) and a weaker correlation between minimum day temperature and ψ_PK_ (r=0.262, p=0.026). This seems to parallel the increased responsiveness to higher temperatures of *per^01^*flies in locomotor activity [47].

### Arrhythmic eclosion under quasi-natural conditions in *per^01^, tim^01^* mutant flies

The results above showed that *per^01^* mutants, but not WT flies eclose arrhythmically under quasi-natural temperate conditions. To verify this effect of an impaired molecular clock, we conducted outdoor eclosion experiments in a subsequent year, using *per^01^; tim^01^* double mutants. Like for *per* [32,48], mutation in the *timeless* (*tim*) gene [34] as well as overexpression of *tim* in the peripheral clock (prothoracic gland) disrupts eclosion rhythmicity [22,23]. Out of the four successful experiments, one was strongly rhythmic (RI=0.44), one was arrhythmic (RI=0.07) and two were just above the border for weak rhythmicity (R=0.12), even though the Zeitgeber amplitudes were comparatively stable during the experiments. Consistent with the results from *per^01^* flies, the eclosion gate was considerably wider compared to WT_CS_ flies (Winfree’s R = 87.7 ± 9.1, a value close to arrhythmicity (R > 90)). The wider eclosion gate and Winfree’s R is statistically similar to *per^01^* mutant flies (Fig. 2E), with *per^01^; tim^01^* flies also showing extended eclosion activity not only into the second half of the night but also into early afternoon. Likely due to stable and strong Zeitgeber amplitudes, *per01;tim01* mutants showed a wild-type like mean eclosion time of 9:34 h ± 5:34 s.d. (n =1918), with a vector length (r) of 0.345 (Fig. 2F).

### Impaired PDF signalling has little effect on eclosion rhythmicity under quasi-natural conditions

The results above suggest that a functional molecular clockwork is required to maintain eclosion rhythmicity under variable temperate outdoor conditions. Next, we monitored eclosion rhythmicity under laboratory and quasi-natural conditions in *pdf^01^ and pdfr (han^530^)^4^*mutant flies which are defective in PDF signalling. PDF is a neuropeptide signal released by a subset of circadian pacemaker cells in the brain (the PDF^+^ small and large lateral ventral neurons (s- and lLN_v_s)) which is received by the PDF receptor PDFR encoded by *han* and expressed by a large set of central clock neurons [37,49–52]. PDF-PDFR signalling is important to maintain stable phase and Ca^2+^ activity relationship between central clock neurons [53]. Besides this clock-internal function, PDF is also a major output factor of the circadian clock [54–57]. We hypothesised that if the molecular clockwork is required for eclosion rhythmicity under natural conditions, then PDF signaling mutants should show rather normal eclosion rhythmicity under quasi-natural conditions, as the individual groups of pacemaker neurons are kept in sequence by the Zeitgebers present. If PDF signalling itself is also required as an output factor, then we expected impaired eclosion rhythmicity.

Under laboratory conditions and consistent with the literature[22,23,48], PDF signalling mutants were rhythmic under LD12:12 (Fig. 1C, D), but became arrhythmic after two to three days in DD (Fig. 1C”, D”) presumably since the individual groups of pacemaker cells lose their proper sequence of activity [23].*pdf^01^* and *han^5304^* mutants also eclosed rhythmically under WC conditions (Fig. 1C’, D’), yet maintained residual rhythmicity under CC (Fig. 1C’”, D’”), albeit with a strongly altered eclosion pattern lacking the clear drop in eclosion events during the second half of the day.

Under quasi-natural conditions, *pdf^01^* and PDFR mutants (*han^5304^*) were mostly rhythmic (67% and 57% respectively, n=7, 8; RI >0.1). The mean eclosion time over all experiments was very similar to WT_CS_; with 09:13 h ± 03:54 s.d. and a vector length (r) of 0.593 for *pdf^01^* flies (n =1918), and 09:44 h ± 05:05 s.d and a vector length (r) of 0.412 for *han^5304^* flies (n =1918). Also Winfree’s R showed strong rhythmicity in 85% of the experiments for *pdf^01^* (R= 43.3 ± 26.4 s.d., n=7) and *han^5304^* (R= 44.3 ± 24.2 s.d., n=8) flies. For both *pdf^01^*and *han^5304^*flies, mean R was significantly smaller (p=0.002, Tukey Multiple comparisons) than for *per^01^* mutants(100.3 ± 17.6 s.d., Fig 2E). A correlation between ψ_PK_ peak eclosion time and maximum/minimum day temperature was found for *han^5304^*PDF receptor mutants (max: r=0.293, p=0.008, min: r=0.601, p<0.001), but was missing in *pdf^01^* flies.

Compiled, these data indicate that natural Zeitgeber can compensate for a loss of PDF signalling (pdf^01^, pdfr, han^530^) but not for the loss of a functional clock (per^01^, tim^01^) in eclosion timing. In other words, our results show that a functional endogenous clock is required for rhythmic eclosion behaviour under variable quasi-natural conditions.

### Humidity as a Zeitgeber is unable to entrain eclosion rhythmicity

Temperature and relative humidity are typically inversely correlated, and it is difficult to separate the influence of these parameters on eclosion timing under quasi-natural conditions. Humidity has been suggested to be involved in eclosion timing of *Drosophila* [43], but there is no direct evidence for this assumption and it is unclear whether humidity can act as a Zeitgeber for *Drosophila*. We therefore tried to entrain CS wildtype flies to humidity cycles of 12 hours 70% and 12 hours 30% relative humidity under constant red light (λ=635 nm) at 20°C throughout the entire development. Under these conditions, flies eclosed arrhythmically (mean RI = 0.09±0.14, N=4, n=664) without distinguishable preference for either the wet or dry phase (Suppl. Fig. 2). This finding shows that, unlike plants [58], *Drosophila* is unable to entrain to humidity cycles and hence does not use humidity as a Zeitgeber.

The timing of eclosion to the morning hours typically is hypothesised to represent an adaptation to higher relative humidity during the morning, reducing water loss and facilitating proper wing unfolding as long as the cuticle is untanned [59]. In fact, the kauri moth, *Agathiphaga vitiensis* requires >80% relative humidity (RH) to successfully eclose (Robinson and Tuck 1976 cf.[60]). We therefore tested whether relative humidity has an effect on eclosion timing or might even prevent eclosion. WT_CS_ and *per^01^* mutant flies were allowed to eclose at normal (60%), high (80%) and extremely low (2%) RH at 25°C, and failure to eclose or to expand the wings upon successful eclosion was noted. Even at 2% RH, ≥ 88% of flies eclosed successfully and expanded their wings (Fig. 2G; WT_CS_: N=6, n=829, *per^01^*: N=6, n=2047). Only for WT_CS_ eclosion was this percentage significantly lower than in both controls at 60 and 80% RH (Fig. 2G, Tukey multiple comparison, p<0.05; WT_CS_: 60% RH: N=6, n=1581; 80% RH: N=8, n=3532; *per*^01^: 60% RH: N=6, n=1936; 80% RH: N=9, n=3340). These results speak against a direct inhibition of eclosion by low RH within natural habitat conditions of *D. melanogaster*, and show that low humidity does not significantly affect wing expansion in the fruit fly. This is against earlier assumptions [42], but in line with results earlier obtained for the much larger onion fly *Delia antiqua* [61].

### Eclosion timing of WT_CS_ but not clock-related mutants is largely unaffected by day-to-day variation in the amplitude and absolute level of abiotic Zeitgebers

To better assess the immediate relation between time of eclosion and abiotic Zeitgebers, and to evaluate eclosion probabilities (0 < p < 1) in response to changes in environmental variables, we next applied a logistic regression model. We included time of day (= hour and hour^2^), temperature, light intensity and nautical dawn as environmental variables, and genotype as a categorial predictor, and ignored interactions of higher than second order. We selected the best statistical model by backward simplification [41], starting with the most complex model. Then, effects of weak significance were removed in a stepwise manner until Anova model comparison yielded a significant difference which indicates that further simplification would result in a substantially inferior model. An example of the actual data for each day and genotype plotted against the best model is shown in Suppl. Fig. 3-4.

All environmental variables contributed significantly (in particular via interaction effects) to the explanation of the observed temporal patterns. In general, eclosion timing is dominated by hour of the day (Fig. 3A, Suppl. Table 1) with an overall tendency of increasing eclosion probabilities over the course of the day but a more or less pronounced peak for eclosion probability before noon (Suppl. Fig. 3-4); this does not exclude that the majority of animals eclose already at an earlier time of the day as the probability that an individual does not eclose before hour h=T is 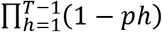. Time of nautical dawn (strictly correlated with date) had a moderate effect with earlier emergence when sun rises earlier (variables were scaled such that early times of dawn have negative values, late times positive values).Whereas temperature and light intensity acted as weaker main factors, both were involved in strong interaction effects with hour of the day (Suppl. Table 1). The quadratic time component (hour^2^), had a highnegative effect strength in WT_CS_, as well as in *per^01^* and *pdf^01^* mutant flies (Fig. 3A, Suppl. Table 1). These negative values indicate a particular peak eclosion per day, which seems to be hardly present in *han^5304^* flies (Fig. 3A). Light, in contrast, had little effect on eclosion except for *per^01^* mutants (Fig. 3B), while temperature had a significant and opposing effect on *pdf^01^* and *han^5304^* flies (Fig. 3C). These results support the finding that eclosion timing in WT_CS_ flies is strongly driven by the endogenous clocks, while the clock and PDF signalling mutants are more susceptible to changes by the main Zeitgeber light and temperature, respectively. Importantly, possession of a functioning molecular clock (WT_CS_, *pdf^01^, han^5304^*) seems to uncouple eclosion behaviour from momentary changes in light intensity. The high effect strength of light on *per^01^* mutants (Fig. 3B) might at least partially explain the significantly earlier eclosion time of *per^01^* mutants (Fig. 2B) with a mean around 6:00, which is inbetween the mean time of nautical dawn and official sunrise during the experimental period (05:04h and 06:23h respectively). The stronger effect of temperature especially on *pdf^01^* mutants is remarkably similar to the stronger temperature dependency of the onset of evening locomotor activity in *pdf^01^* mutants under LD conditons [62]. For locomotor activity, the PDF-expressing sLNvs are a central part of a light-entrainable oscillator that dampens the temperature effect on the phase of the clock neuron subsets (dorsal neurons (DNs) and lateral posterior protocerebrum neurons (LPNs)) constituting the temperature-entrainable oscillator [62]. Hence also for eclosion, lack of PDF signalling that establishes phase relationship between the different oscillators [53] may lead to a stronger contribution of the temperature-entrainable oscillator during eclosion timing. This implies also a significant effect of temperature on PDFR mutants (*han^5304^*) which indeed is the case (Fig. 3, Suppl. Table 1). It is, however, difficult to explain why *pdf^01^*flies tend to eclose at cooler temperatures than WT_CS_, while *han^5304^*flies tend to eclose at warmer temperatures. Interestingly, a similar discrepancy was found in the temperature preference, which is towards lower temperature in *pdf^01^*flies, but wildtype-like in *han^5304^* mutants [63]. Altered temperature preference may thus also contribute to the increased negative effect strength of temperature on *pdf^01^* mutants.

**Fig 3:**
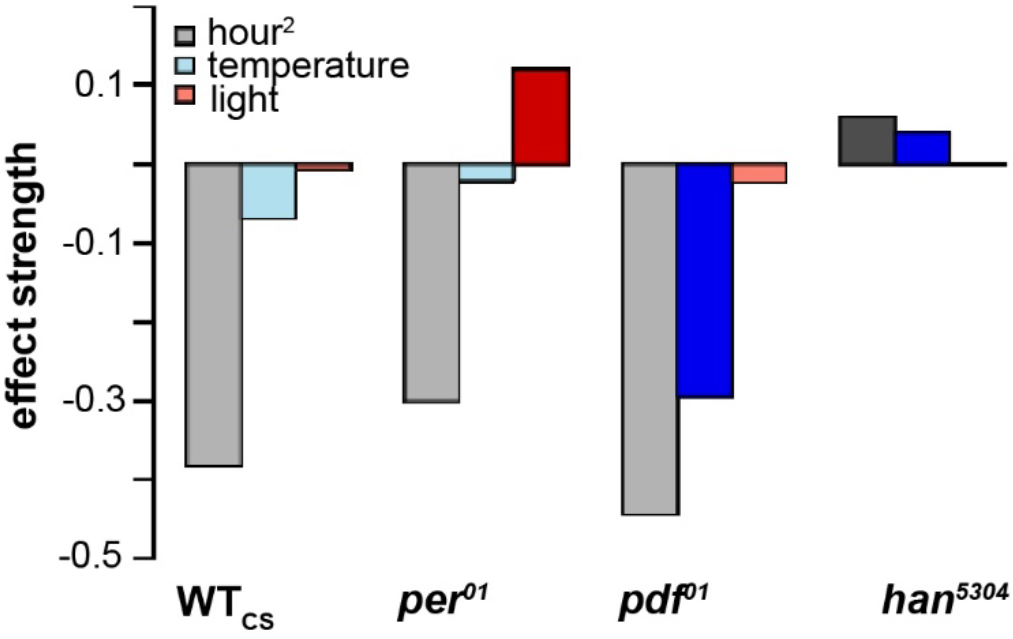
Effect size for effects that show a significant interaction effect with the genotype. Independent variables scaled to mean; hour is scaled to 12:00=0. Darker colors indicate significant differences to the WT_CS_reference. Note that effect strengths cannot directly be compared between factors.

## Conclusion

Our results show that a functional molecular clock is required for normal eclosion rhythmicity under quasi-natural temperate conditions in the fruit fly *Drosophila*. This is in contrast to the results on locomotor activity in *per* mutant mice and fruit flies under comparable conditions. While a general circadian dominance of behaviour has recently be questioned [13], our results demonstrate that circadian dominance cannot generally be excluded, and is prevailing at least for the daily timing of *Drosophila*. Clear latitudinal and altitudinal differences in eclosion rhythms have been found for various *Drosophila* species [64–67], making it reasonable to assume that eclosion timing serves an adaptive function and hence implying a significant adaptive value to the circadian clock under natural conditions. The ultimate causes that time eclosion to a particular time of the day in many if not most insects nevertheless still remain to be identified.

*Drosophila melanogaster* originated in the tropical Africa south of the Sahara, colonized Europe and Asia about 15,000 years ago, and was brought to the Americas and Australia some hundred years ago [68,69]. As cosmopolitan species, *D. melanogaster* adapted to many different climates, ranging from tropical regions with rather stable day-to-day conditions to temperate regions with more variable climatic conditions. This adaption lead to latitudinal clines of different morphological and behavioural traits [69] including different courtship and mating behavior [70]. It is therefore interesting to note that *per^01^* mutant flies under seminatural tropical conditions in Bengaluru, India eclosed rhythmically, with a rhythmicity stronger than under light:dark conditions in the laboratory [43]. This is not contradicting our findings, as the prevailing weather conditions are much more stable in Bengaluru, with daily temperature curves more predictable and regular and much reduced variability in day-to-day light intensity and temperature amplitudes as compared to the temperate conditions around Würzburg. Such stable conditions were rarely met during our experiments under temperate conditions, but when such conditions prevailed they also produced rhythmic eclosion in *per^01^* mutants. Thus, the difference of eclosion rhythmicity and robustness between temperate and tropical conditions might in large part reflect differences in temperature cycles and their masking effects.

Moreover, and albeit the observed differences in rhythmicity, eclosion gate width increased and the eclosion phase shifted towards the night in *per^01^* flies under tropical [43] similar to what we found for temperate conditions. It is therefore tempting to speculate that the circadian dominance of eclosion is present but of lesser importance in the tropics. With the spread of *D. melanogaster* into subtropical and temperate regions worldwide, circadian dominance gained in importance and ensures rhythmic eclosion at the right time of the day also at higher lattitudes.

## Data accession

The original data will be archived at Dryad repository (datadryad.org).

## Authors’ contributions

FR and CW designed the study, FR and STM carried out the eclosion experiments, MH carried out the humidity experiments, CW and DR supervised the practical study, OM and TH developed the statistical model, FR, OM, TH, STM, MH, DR and CW analysed the data, CW and FR wrote the manuscript draft with help from OM, TH and DR. All authors read and approved the manuscript

## Acknowledgments

We thank the University’s Bee station (Ricarda Scheiner, Dirk Ahrens-Lagast) for providing space for the outdoor shelter, Heike Wecklein, Konrad Öchsner, Johann Kaderschabek and Susanne Klühspies for excellent technical assistance, John Ewer for sharing the Matlab scripts, and Koustubh Vaze for inspiring discussions and critical reading of the manuscript.

