## Supplementary Material for "Natural Zeitgebers cannot compensate for the loss of a functional circadian clock in timing of a vital behaviour in *Drosophila*"

**Supplementary Table 1**. Summary of parameter estimates for the best fitting model. First column specifies estimated model parameters for the reference genotye CS±s.e.m. (significant values plotted in bold), the other values in the genotype colums significant differences in parameter estimates for the other genoytpes. The last column lists significant interaction terms between the environmental covariates.

| $WT_{CS}$ | | $per^{01}$ | $pdf^{01}$ | $han^{5304}$ | Interactions between environmental variables |
|---|---|---|---|---|---|
| **-1.29±.1** | intercept | -- | -- | -0.56 | |
| **0.55±.03** | hr | -- | -- | -- | temperature (-.31***), light (-.53***), dawn (-.28***) |
| **-0.38±.07** | hr^2 | -- | -- | 0.44 | temperature (.15***), light (-.18**) |
| -0.07±.05 | temperature | -- | -0.23 | 0.11 | light (.07**) |
| 0.01±.04 | light | 0.12 | -- | -- | dawn (-.13***) |
| **-0.14±.06** | dawn | -- | -- | -- | |

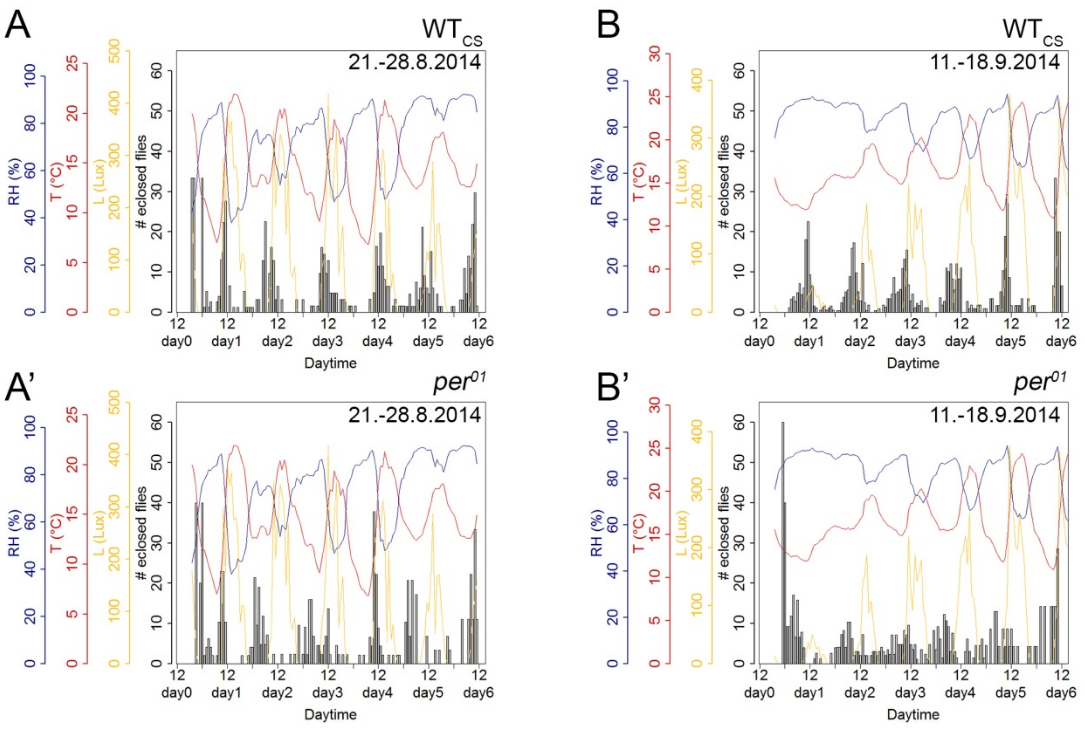


***Supplementary Fig 1***: *Original eclosion profiles of* $WT_{CS}$ *(top row, A-B) and* $per^{01}$*mutant flies (bottom row, A'-B') for two different weeks end of August (A-A') and mid of Septembre (B-B'). Relative humidity (RH) is shown in blue, Temperature (T) in red, and light intensity (L) in yellow.*

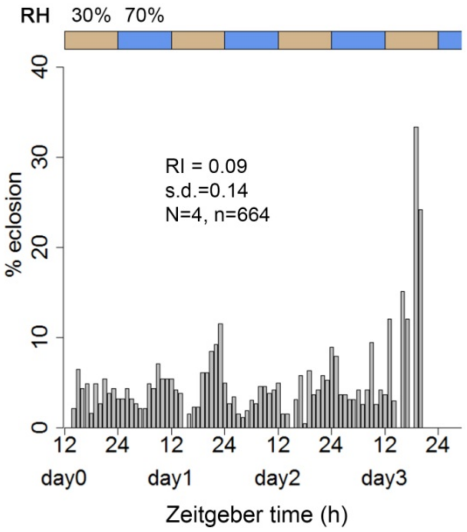


***Supplementary Fig 2***: $WT_{CS}$ *flies were kept in DD and a humidity cycle of 12h 30% and 12h 70% RH was applied. Despite the humidity cycle, flies eclosed arrhythmically, indicating that humidity changes are unable to entrain eclosion rhythmicity.*

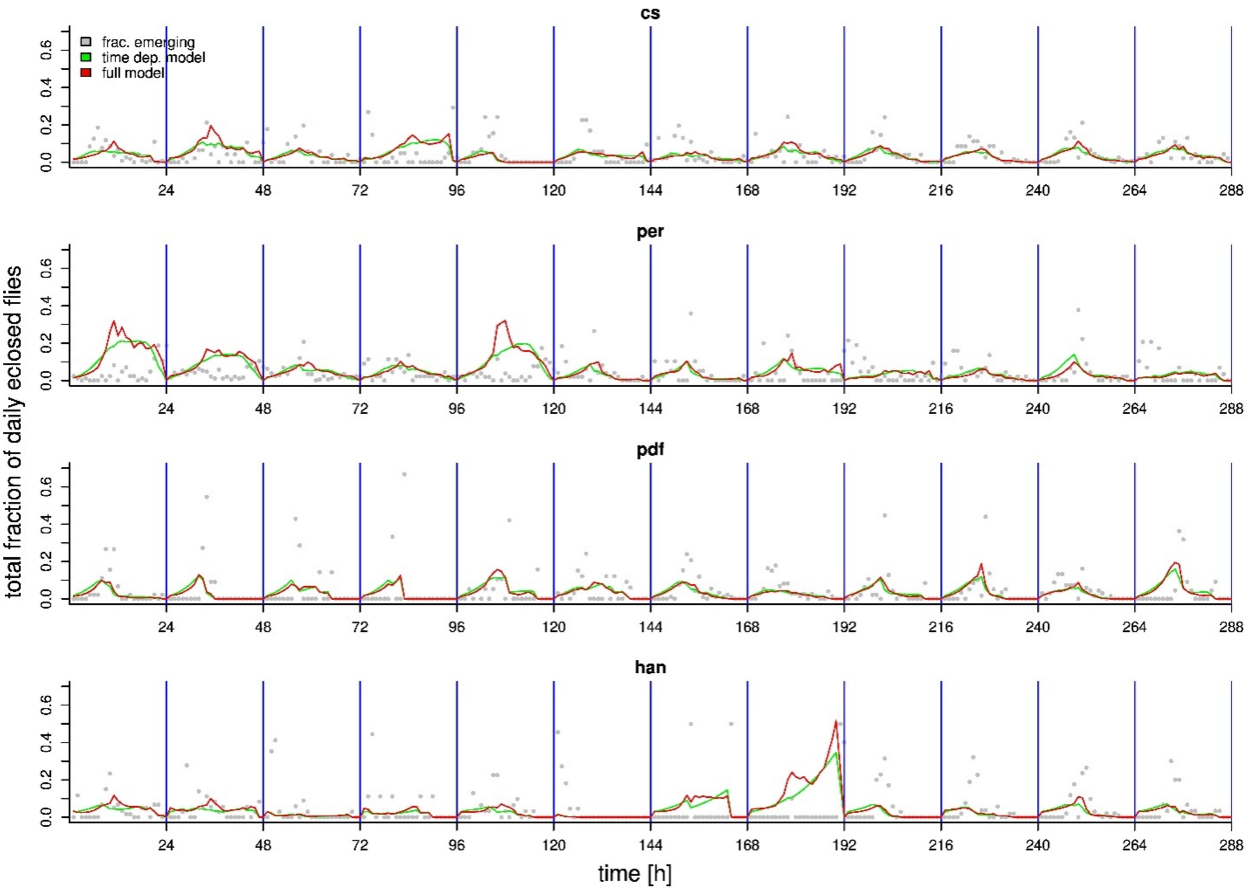


**Supplementary Fig 3.** Comparison of model predictions between four genotypes. Grey points shows the fraction/hour of flies eclosing during a day. The green line shows the predicted emergence based on hour and hour$^2$ only whereas the red line shows the curve for the best fitting model also accounting for the effects of temperature, light and time of sun-rise (dawn). For clarity the figure shows only data and model for 12 days.

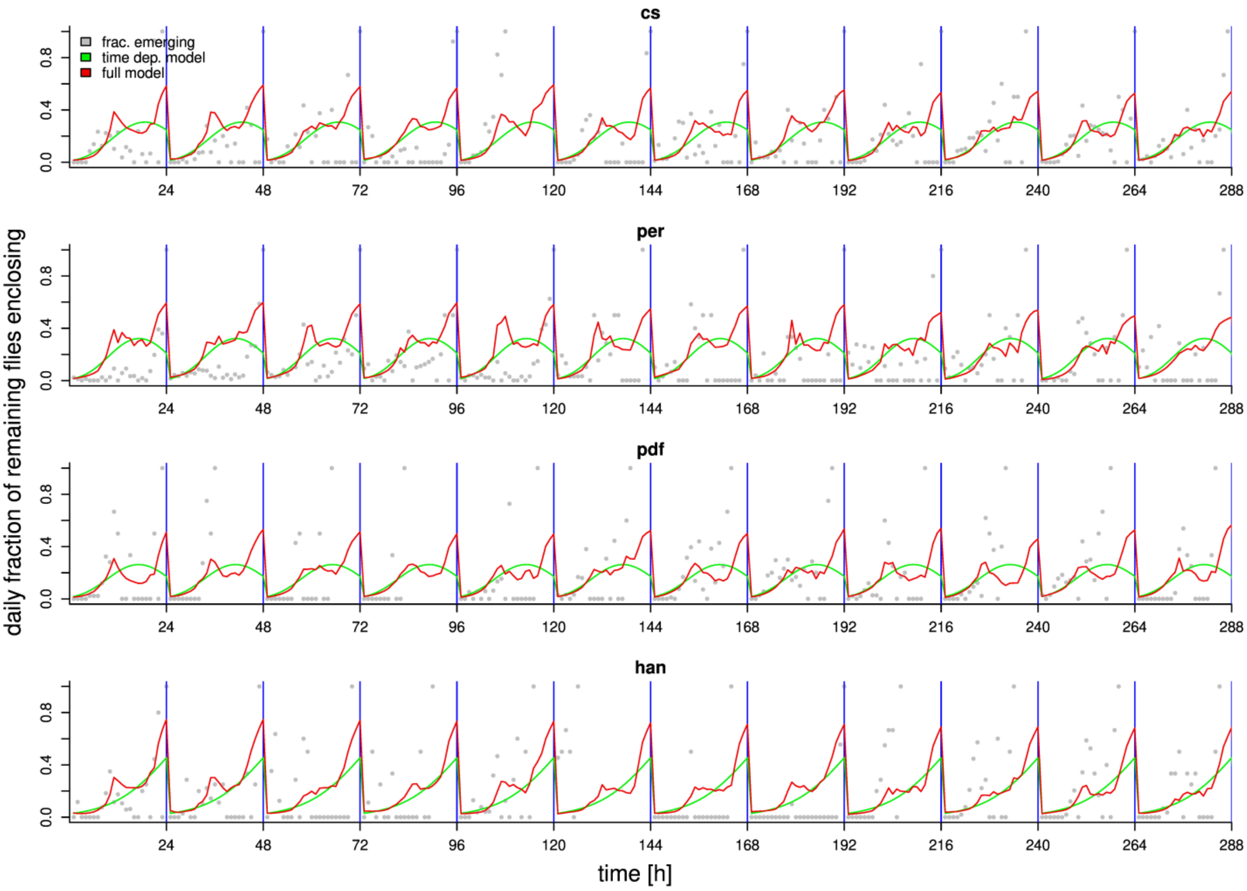


**Supplementary Fig 4.** Comparison of model predictions between four genotypes. Grey points shows the proportion of flies eclosing during each hour that did not emerge up to this moment during that day. Note that the last empirical observation during the day must thus take the value 1 when the last flies of the day emerge. The green line shows the predicted emergence based on hour and hour² only whereas the red line shows the curve for the best fitting model also accounting for the effects of temperature, light and time of sun-rise (dawn). For clarity the figure shows only data and model for 12 days.
